# Spoti-find: A novel, open-source void spot assay image analysis tool

**DOI:** 10.1101/2025.09.30.679176

**Authors:** Cara C. Hardy, Susan Sheehan, Seamus Mawe, Nick Sebasco, Will Ricke, Zaenab Dhari, Stephen J. Crocker, Matt Mahoney, Ron Korstanje

**Affiliations:** The Jackson Laboratory, Bar Harbor, ME, USA; Department of Urology, School of Medicine and Public Health, University of Wisconsin-Madison, Madison, WI, 53705, USA; Department of Neuroscience, University of Connecticut School of Medicine, Farmington, CT 06032, USA; Department of Immunology, University of Connecticut School of Medicine, Farmington, CT 06032, USA

## Abstract

The void spot assay (VSA) is a widely used, non-invasive method for evaluating urinary behavior in rodents, but existing analysis tools are limited in scope, throughput, or accessibility. We developed Spoti-find, a stand-alone, open-source VSA image analysis application that introduces novel, biologically meaningful metrics including void circularity, distance to paper edge, and volume-based binning into primary, micro-, and nanovoids. Designed with usability in mind, Spoti-find features a graphical interface that enables manual or semi-automated spot identification, adjustable thresholds, and streamlined data export without the need for coding expertise. We validated Spoti-find across diverse datasets, showing strong inter-user consistency, sensitivity to known phenotypes in aging and disease models, and accuracy in capturing novel parameters while demonstrating high agreement with existing tools. By capturing behavioral context and spatial morphology in voiding patterns, Spoti-find expands the interpretive power of VSA and provides a flexible, user-friendly platform for phenotyping urinary dysfunction in preclinical studies.

---

Urinary dysfunction is a hallmark of numerous neurological, metabolic, and age-related diseases, yet the biological basis of aberrant voiding behavior remains difficult to study in preclinical models. The void spot assay (VSA) provides a non-invasive, low-cost method to assess urinary behavior in rodents by capturing the number, size, and distribution of urine spots deposited on filter paper over time (Hill, Zeidel et al. 2018). Its simplicity, flexibility, and relevance to multiple organ systems -- including the bladder, kidney, central nervous system, and musculoskeletal system -- make it a powerful tool for investigating urinary phenotypes across diverse experimental contexts.

The most widely used platform and current gold standard, Void Whizzard, was a pivotal advancement for the field. By offering semi-automated analysis of VSA images within ImageJ, it made standardized spot detection, sizing, and classification broadly accessible for the first time (Wegner, Abler et al. 2018). More recently, ML-UrineQuant extended these capabilities by incorporating deep learning for high-throughput analysis, enabling automated spot classification across large datasets with adaptable parameters tailored to specific lab needs (Wegner, Abler et al. 2018). Despite these advances, both platforms have limitations: they are limited in capturing novel biologically relevant metrics with clinical implications, often require extensive troubleshooting and quality control (QC) checks by the experimenter, and in the case of machine learning–based approaches, the opaque ‘black box’ nature of the algorithms can make resolving issues nearly impossible.

To address the need for biologically meaningful VSA analysis that remains accessible and easy to use, we developed *Spoti-find*, a stand-alone, open-source Python-based application. Spoti-find builds upon prior tools by introducing analytical parameters that capture the spatial and morphological context of voiding behavior, features that may reflect underlying physiological or behavioral states. These include: (i) <u>void circularity</u>, a proposed proxy for intentional voiding versus passive leakage; (ii) <u>distance from the edge of the filter paper</u>, which may reflect anxiety or urgency-like phenotypes based on rodents’ preference for voiding along boundaries; and (iii) <u>void size binning</u>, which classifies spots into primary voids, microvoids, and nanovoids using biologically interpretable volume cutoffs. Designed with usability in mind, Spoti-find features an intuitive graphical interface, allows both automated (rectangle) and manual (polygon) spot selection, enables adjustable thresholds, and streamlines data export. While different platforms may better suit different experimental goals—such as ultra-high throughput versus detailed spatial metrics -- Spoti-find offers a customizable and behaviorally informative alternative that expands the analytical utility of VSA for diverse preclinical applications.

Importantly, Spoti-find was designed for accessibility and transparency. The tool requires no coding knowledge, features a clean graphical interface, and allows adjustable thresholds for all void classification parameters. While Spoti-find is a manual program and does not prioritize high throughput, it enables rigorous and reproducible data extraction with minimal QC effort, even across variable experimental conditions. In this study, we validate Spoti-find across multiple independent datasets and demonstrate its ability to detect known disease- and age-associated urinary phenotypes, while uncovering additional spatial and morphological features that may offer translational insights. By emphasizing interpretability, flexibility, and biological relevance, Spoti-find represents a next-generation approach to VSA analysis that bridges the gap between simple descriptive metrics and high-dimensional behavioral phenotyping.

## Results

### Spoti-find captures novel voiding parameters with biological relevance

To expand the analytical utility of the VSA, Spoti-find introduces several new metrics designed to capture spatial and morphological features of voiding behavior. The user interface allows experimenters to adjust specific parameters, select spots using either a rectangle tool that identifies the spots automatically or a polygon tool that allows users to manually outline and identify spots using the fill function (Fig.1a). Users are also able to calibrate volume detection using spots of known sizes (*User Manual; see Supplemental Information*). Spot binning was implemented to distinguish between primary voids (>22 µL), microvoids (2–22 µL), and nanovoids (1–2 µL) (Fig.1b), based on prior cystometric studies and published estimates of typical voiding volumes (Yu, Ackert-Bicknell et al. 2014, Kim and Hill 2017, Wegner, Abler et al. 2018, Fraser, Smith et al. 2020). Spots under 1 µL were classified as noise but remain accessible to users analyzing disease models where smaller volumes may be biologically meaningful (e.g., benign prostatic hyperplasia, BPH (Lee, Yang et al. 2015). Additionally, our tool introduces a circularity metric, reported as a value between 0 and 1, where 1 denotes a perfect circle and 0 reflects a straight line. This metric enables identification of ‘worm-like’ elongated voids—presumed to occur while the animal is in motion—versus rounder, more stationary voids (Fig.1c).

**Fig. 1.**
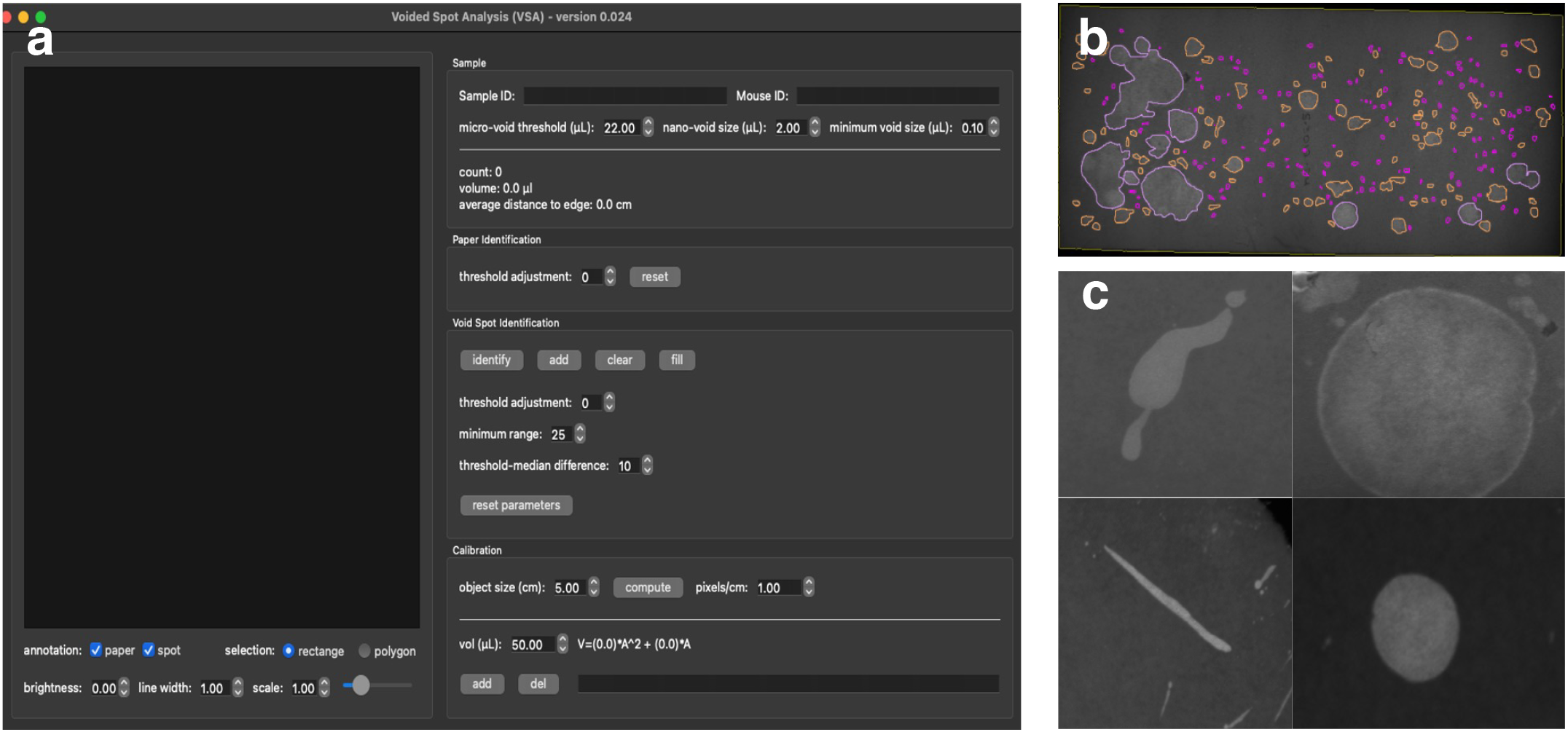
Spoti-find enables quantification of novel spatial and morphological features in void spot assay analysis. a) Graphical user interface for Spoti-find is easy to use and adjustable to suit specific experimental parameters. b) VSA image denoting primary (light purple), micro (orange), and nano voids (pink). c) Elongated “worm-like” voids and circular voids.

Mice exhibit a strong preference for voiding near the edges of their environment, a behavior thought to be evolutionarily adaptive by helping them avoid open, exposed spaces, a phenotype consistent with their natural thigmotaxic tendencies (Yu, Ackert-Bicknell et al. 2014). Increased center voiding due to increased time spent in this sector has been hypothesized as a potential indicator of anxiety-related phenotypes (Biallosterski, Prickaerts et al. 2015). Therefore, we sought to capture the spatial location of each void relative to the paper edge, using the shortest linear distance from the center of each spot to the nearest boundary. This measurement provides a novel proxy for spatial behavioral tendencies in rodent models, leveraging the observation that healthy mice preferentially void along the cage perimeter. Increased central voiding may indicate altered urgency, incontinence, or anxiety-like phenotypes. These novel parameters enable users to extract new behavioral insights from VSA data, without requiring complex computational tools or video tracking systems. By making biologically relevant features like spot shape and location easily quantifiable, Spoti-find enhances the interpretive power of the assay while preserving usability and transparency.

### Spoti-find has strong agreement with existing analysis tools and between users

To assess whether Spoti-find performs comparably to existing VSA tools, we evaluated agreement between Spoti-find and the current gold standard, Void Whizzard, for core metrics including total spot count. Although Void Whizzard bins voids by diameter and focuses primarily on spot number rather than biologically informed categories, we found strong correlation (R^2^= 0.9161737 p=2.2x10^−16^) between tools for spot count between images analyzed in Void Whizzard and Spoti-find (Fig.2a). Notably, the Void Whizzard analyses were conducted at different institutions by researchers with expertise in VSA methodology, using datasets collected under diverse experimental conditions and timeframes. Spoti-find analyses were subsequently performed on the same raw images by users without specific training in urologic research, demonstrating that the tool enables accurate and unbiased analysis independent of prior domain experience. The strong agreement between Spoti-find and Void Whizzard across these varied datasets highlights the accessibility, ease of use, and analytical reliability of Spoti-find, even when used by non-expert operators. This robustness across institutions, users, and timepoints reinforces Spoti-find’s utility as a widely adoptable platform for VSA analysis.

**Fig. 2.**
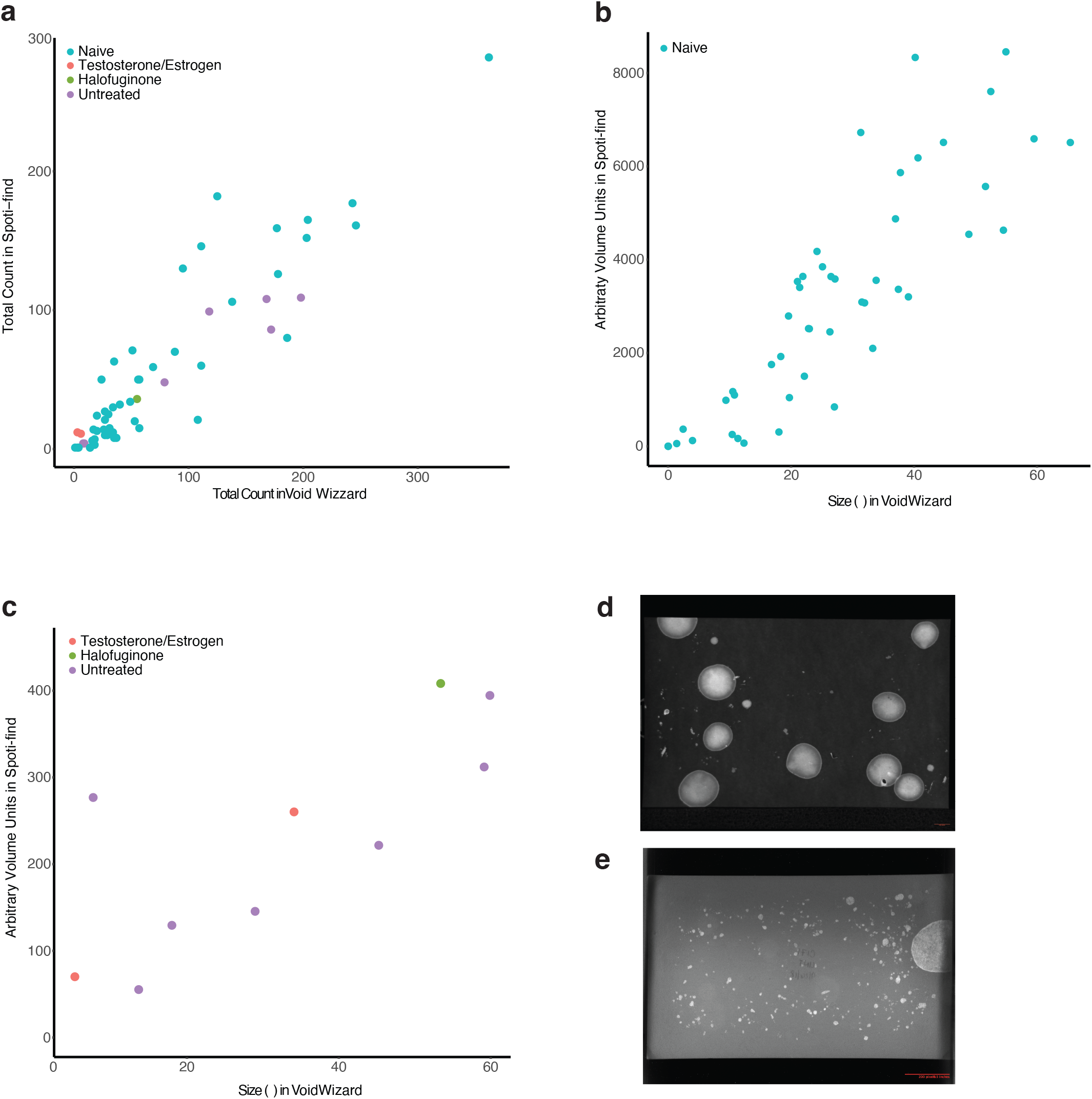
Validation of Spoti-find performance against existing tools and across users. **a**. Spoti-find and Void Whizzard show high agreement with for total number of spots regardless of experimental conditions. Spot sizes/volumes are highly correlated between Spoti-find and Void Whizzard in both naïve animals (**b**) and treated animals (**c**), however variation between experimental conditions, imaging settings, lightings, and paper used complicate the ability to compare between different studies. Examples of different paper types, imaging conditions, and spot patterns are shown in **d**) and **e**).

Correlations between spot volumes in Void Whizzard and Spoti-find were high for both naïve animals of different ages (R^2^ =0.8643384, p=2.495x10^−15^) (Fig.2b) and animals of the same age under different treatment conditions (R^2^=0.7805848, p=0.007707.) (Fig. 2c). However, despite strong agreement in spot counts, correlations in volume were less robust when comparing across experimental conditions (R^2^= 0.6233786, p=1.726x10^−7^). This discrepancy likely stems from variability in imaging factors such as lighting, camera settings, and paper types, highlighting the need for caution when comparing historical datasets collected under heterogeneous conditions (Fig. 2D–E). For example, in Fig.2b an approximately 40 uL void corresponds to ∼4000 arbitrary volume units (AVUs), whereas under different imaging conditions (Fig.2c) a void of similar size corresponds to only ∼300 AVUs. To account for such inter-experimental variability, all Spoti-find volume measurements are reported in AVUs, since datasets were acquired across multiple laboratories using diverse imaging hardware, paper types, and acquisition parameters. These differences preclude the use of a universal calibration curve to convert pixel-based measures into absolute microliter values. Instead, Spoti-find estimates volume as a function of spot area and pixel intensity, both derived from pixel data and thus highly sensitive to acquisition settings. While spot counts are robust to these variables (Fig.2a), volume comparisons across datasets lacking consistent imaging conditions should be interpreted in relative rather than absolute terms (Fig.2b-e).

To determine the practical robustness of Spoti-find for large-scale dataset analysis, we tested whether user fatigue affected performance across prolonged analysis sessions. Individual users were asked to analyze batches of 10, 20, 30, or 40 papers without taking breaks. Across multiple trials, we found that users were consistently able to complete up to 30 papers per session before experiencing accuracy declines or noticeable fatigue, whereas 40 papers proved unsustainable. We note that this threshold may be model dependent; for instance, diabetic mouse models tend to produce more frequent, smaller voids that require more manual input than models in which voiding is dominated by fewer large-volume primary voids.

We also evaluated inter-user agreement to assess how user variability may impact Spoti-find results. Most parameters showed high concordance across users (Supplementary Table 1), demonstrating that the tool is generally robust to individual analytical styles. However, variability was higher in parameters like circularity and average nano/micro void volume, which are more sensitive to minor differences in polygon selection. For example, small linear segments introduced while tracing void perimeters can disproportionately affect circularity in small spots due to their lower overall circumference. These findings underscore the importance of consistent technique and cautious interpretation when drawing conclusions from highly granular parameters.

Finally, we compared outcomes between Spoti-find and Void Whizzard for previously analyzed datasets modeling multiple sclerosis (cuprizone model) and age-related voiding phenotypes. Across both aging (Fig.3) and disease models (Fig.4), Spoti-find successfully recapitulated known group differences in standard VSA metrics, reinforcing its reliability for biologically meaningful inference.

**Fig. 3.**
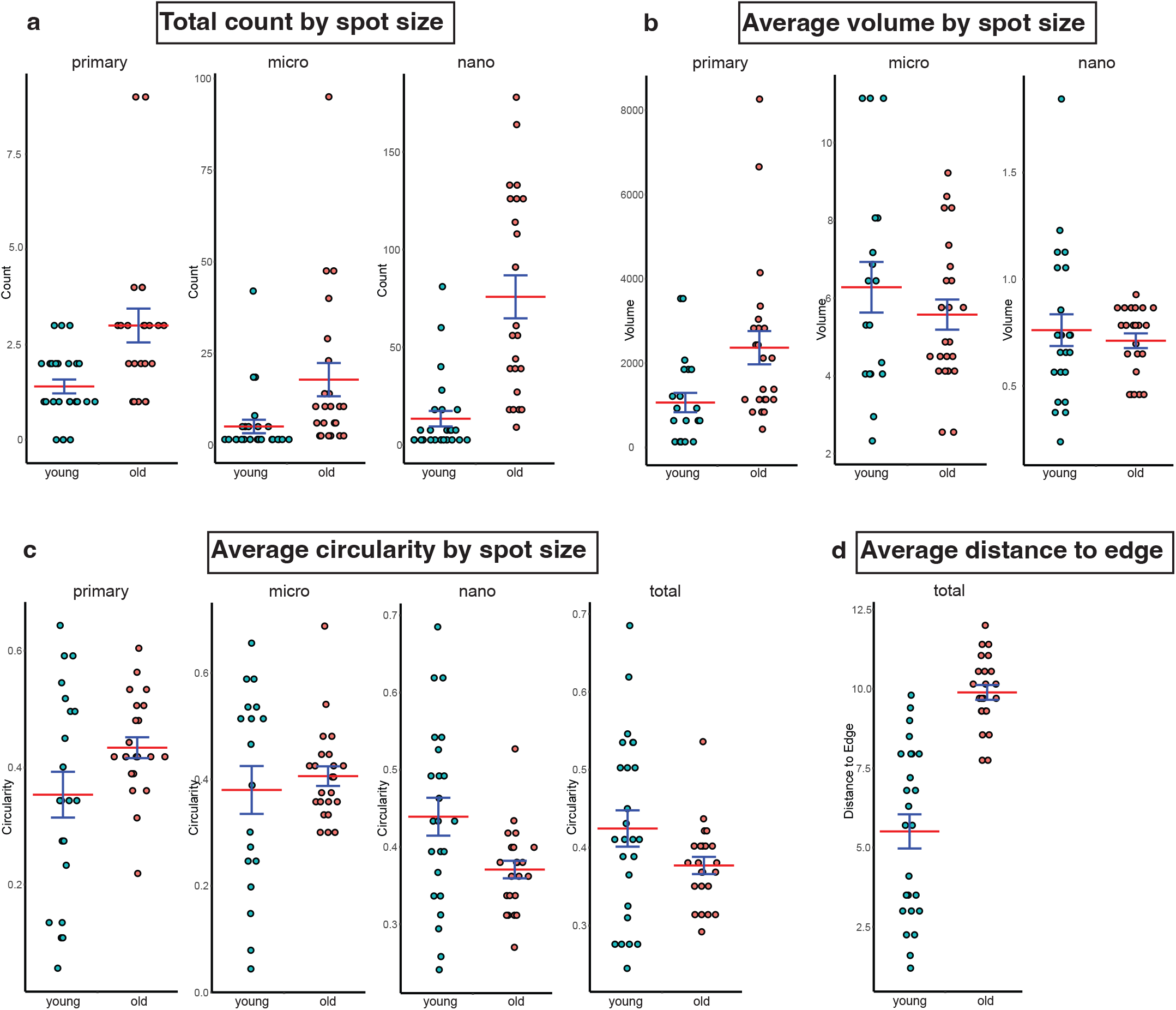
Spoti-find detects voiding behavior differences attributable to aging. Comparison of void counts by spot size (**a**) and volume by spot size (**b**) between young and old mice. **c**) Average total circularity and circularity by spot size between young and old mice. **d**) Average spot distance to paper edge between young and old mice.

**Fig. 4.**
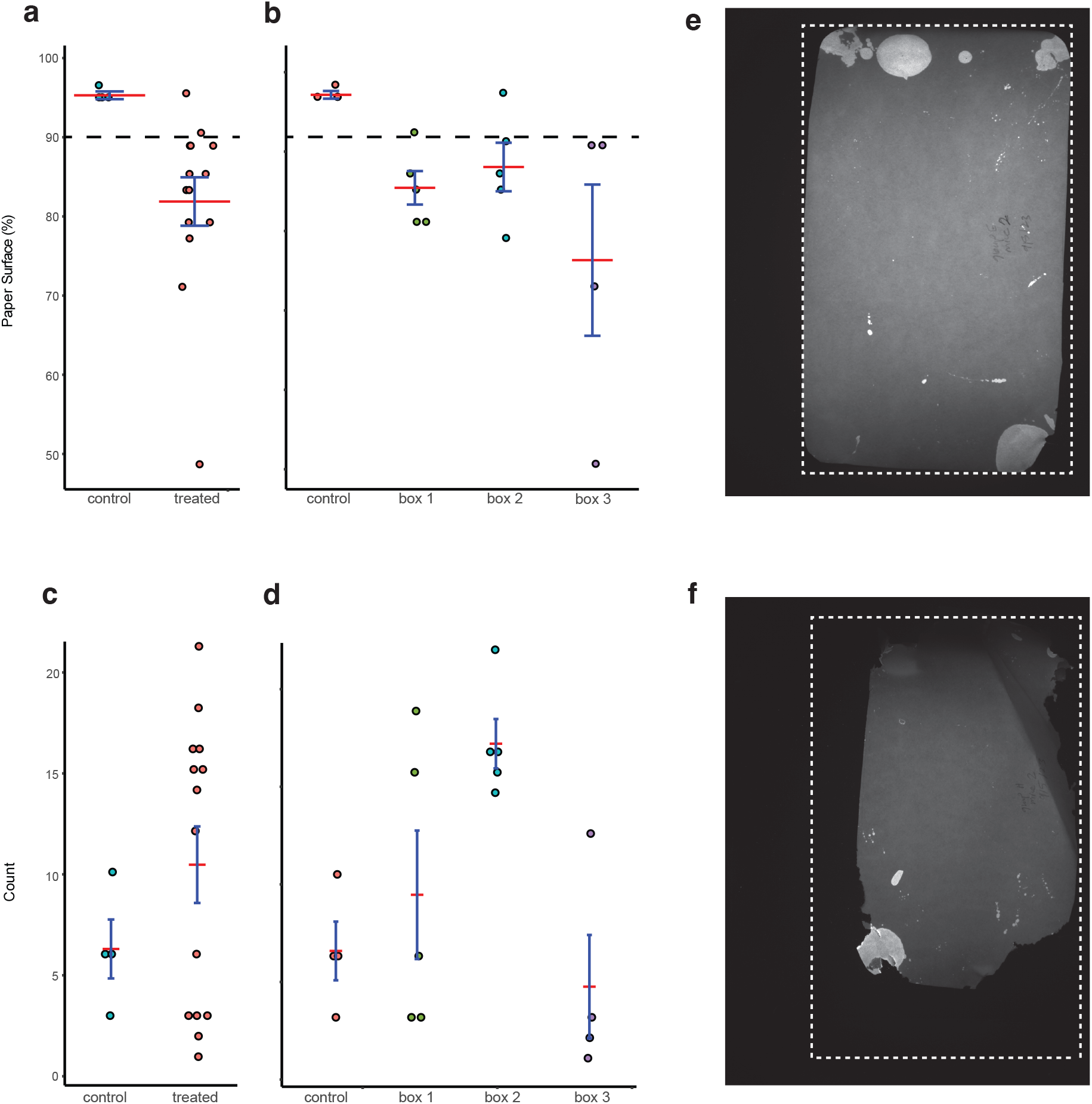
Experimental design considerations to improve the accuracy and translatability of VSA analysis using a cuprizone model of demyelination. **a**) Paper area differs significantly between groups due to destruction by mice. **b**) Cage effects on paper eaten are observed in cuprizone treated mice. **c**) Cuprizone treated mice exhibit higher variability in spot count, with observed cage effects (**d**). **e**) Representative image of a VSA from control (untreated) mice. **f**) Representative image of a VSA from cuprizone treated mice. Dashed lines indicate the standard paper size, demonstrating the significant extent of paper loss shown in (**f**).

### Novel and known age-related alterations in voiding behavior and clinically relevant aging features are captured by Spoti-find analysis

Urinary dysfunction (including increased frequency, reduced bladder capacity, and high post-void residual volume) is a well-documented consequence of aging in both humans and rodent models and behavioral assays like the VSA have proven sensitive to these changes in strain- and age-dependent studies (Yu, Ackert-Bicknell et al. 2014, Kim and Hill 2017, Hill, Zeidel et al. 2018, Bartolone, Ward et al. 2020). Additionally, bladder diaries, which record frequency, volume, and voiding patterns over several days (Mehta, Geng et al. 2023), remain a first-line clinical tool for assessing urinary dysfunction in older adults. They routinely capture hallmark age-related changes such as increased frequency of small-volume voids, larger but less frequent primary voids, and episodes of urgency or incontinence. The VSA serves as a direct preclinical counterpart to this assessment, and Spoti-find allows us to extract features that closely align with diary-based readouts.

Aged C57BL/6J mice exhibited greater variability in spot counts across all size bins, with the most pronounced differences observed in the nano range (primary p=0.00129, micro p=0.00993, nano =1.57x10^−6^) (Fig.3a). These patterns reflect the emergence of small-volume spotting events that are characteristic of aged models. Unsurprisingly, aged animals displayed higher primary void volumes relative to young controls, consistent with their larger body size and bladder capacity at this age, while micro and nano void volumes were comparable between groups (primary p=0.0808, micro p=0.336, nano p=0.548) (Fig.3b), likely due to the narrow absolute range of these bins. Circularity analysis provided a proxy for voiding intention: primary voids were more circular in aged animals, no differences were observed in the micro bin, and nano voids were less circular in aged animals (primary p=0.07067, micro p=0.6451, nano p=0.06284, total p=0.1744) (Fig.3c). This suggests that while primary voiding behavior remains intact with age, small-volume voids are less intentional in nature. When averaged across all size bins, aged animals exhibited an overall reduction in circularity (Fig.3c), supporting the interpretation that Spoti-find can detect incontinence-like phenotypes previously described in these models. Notably, this pattern is consistent with cystometric findings showing that aged animals can perform complete primary voids but also exhibit additional low-volume leak events, sometimes reported as a “dysfunctional responder” phenotype. Finally, the average distance to the paper edge was greater in aged animals compared to young controls(p=2.132x10^−10^) (Fig.3d), further highlighting the maladaptive voiding behaviors that arise with age. Collectively, these data demonstrate that Spoti-find captures both expected and novel age-associated changes in voiding behavior across multiple analytical dimensions.

### Spoti-find captures expected differences between control and cuprizone treated animals

Neurodegenerative diseases frequently impact lower urinary tract function, with symptoms such as urgency, frequency, and incontinence reported across conditions including Parkinson’s disease, Alzheimer’s disease, and multiple sclerosis. These changes are often driven by disruptions in central or peripheral neural control of the bladder, making voiding behavior a valuable functional readout in preclinical models. Multiple sclerosis (MS) is well established to cause neurogenic bladder symptoms in both patients and animal models, including increased voiding frequency, detrusor overactivity, and impaired voluntary control (Ponglowhapan, Church et al. 2008). Because these phenotypes may not always manifest as changes in total urine volume or spot number alone, novel spatial and morphological parameters—such as circularity or defining edge-oriented voiding behavior—may offer additional insight into disease severity or progression. By enabling these metrics to be quantified, Spoti-find provides a new lens through which to assess urinary dysfunction in models of neurologic disease.

To this end, we applied Spoti-find to the cuprizone model of MS-associated demyelination to test whether expected voiding differences could be captured. While group-level differences were observed, these experiments also underscored the importance of experimental context and rigorous QC in interpreting VSA data. Cuprizone-treated mice showed a pronounced tendency to consume the filter paper used in assays(p=0.0564) (Fig.4a), often reducing intact paper area below our 90% QC cutoff for reliable analysis. This variability reflects both the nature of the model, since cuprizone is delivered as a food additive, and intracage dynamics, where social hierarchies can influence individual exposure and, in turn, the degree of demyelination. Indeed, when animals were examined by cage, distinct cage effects emerged (Fig.4b and d), which also contributed to variability in total spot counts (Fig.4c-d). Reduced analyzable area further exacerbated this effect, as spot counts appeared lower simply because urine spots were lost to paper consumption, a phenomenon previously noted in other models such as diabetes. Paper loss also distorted spatial metrics: in cases where large portions of the filter were removed, remaining voids were artificially scored as being extremely close to the edge (Fig.4e-f). These findings highlight the necessity of stringent QC measures and careful interpretation of VSA data, where behavioral artifacts can obscure true biological differences.

## Discussion

Spoti-find represents a next step in the evolution of VSA analysis, building on the success of Void Whizzard, which brought structure, reproducibility, and accessibility to the field. By introducing new biologically relevant parameters such as circularity, void size binning, and distance from the paper edge, Spoti-find expands the scope of insight that can be extracted from this inexpensive and non-invasive assay. While traditional metrics like total spot count and voided volume remain valuable, they often overlook the behavioral (Chen, Zhang et al. 2017) and spatial (Hill, Zeidel et al. 2018) nuances of voiding that may be particularly relevant in models of aging, neurodegeneration, or systemic disease. Spoti-find offers a path toward integrating these dimensions, enabling deeper phenotyping and improved translatability.

Importantly, we see Spoti-find not just as a new tool, but to make more of the data we already collect. The VSA is uniquely suited to longitudinal, non-terminal assessment of urinary behavior in rodents. It can be performed repeatedly, even across the full lifespan of an animal, with minimal handling and without the need for surgical instrumentation like assessments of urinary physiology directly. This makes it especially powerful in models where animals are scarce or valuable, such as neurodegenerative disease models or lifespan cohorts, and aligns with the broader scientific goal of reducing animal numbers while increasing data yield per subject. Because Spoti-find makes it easier to quantify more parameters from each assay, it extends the utility of existing datasets and improves the granularity of behavioral phenotyping without increasing experimental burden.

We also highlight several critical experimental insights enabled by Spoti-find. Acclimation time was found to be of particular importance, as mice, regardless of sex, require at least two full days in their housing cages before voiding behavior stabilizes. This finding underscores the need to account for behavioral adaptation when designing VSA studies and suggests that early time points may not reflect true baseline phenotypes. To maximize the biological value of VSAs, it is critical not only to refine how data are collected, but also how they are analyzed. Spoti-find supports output formats compatible with more rigorous statistical approaches, including linear mixed models (LMMs), which allow for greater sensitivity and flexibility compared to traditional summary statistics. The typical method of averaging across replicates and then averaging across animals (commonly referred to as the “average of averages” approach) risks obscuring biologically relevant variability and inflating false significance due to inappropriate aggregation. In contrast, LMMs retain within-subject variance across repeated measures, account for random effects (such as cage or user), and more accurately reflect the hierarchical structure of preclinical behavioral data. These advantages are particularly relevant for urinary phenotyping, where spot number and distribution can vary meaningfully across time and individuals. This statistical improvement, when paired with Spoti-find’s detailed output structure, allows researchers to make more rigorous, nuanced inferences from their data, an essential step for behavioral studies with subtle or distributed phenotypes.

Although we were not yet able to implement automatic identification of overlapping voids (such as overlapping spots or “spots within spots”) (Wegner, Abler et al. 2018), which human observers can easily discern, this remains a high-priority area for future development. Incorporating this feature would increase the accuracy of voided volume and bin classification, particularly in models with high voiding frequency. Full automation of VSA analysis remains a difficult challenge due to variability in paper type, lighting conditions, spot contrast, and image quality between datasets and experiments performed at different institutions. For these reasons, Spoti-find was built to balance automation with user control. The polygon tool allows for precise manual correction where automation falls short, and adjustable thresholds support flexible application across experimental contexts.

One of the most exciting aspects of Spoti-find is its ability to approximate behavioral features that previously required costly or low-throughput methods like metabolic cages or video tracking. Intentional voiding behavior— such as stopping, squatting, and releasing a coordinated void—is difficult to detect without invasive or intensive monitoring setups (Keller, Chen et al. 2018). By introducing metrics like circularity and spatial distribution, Spoti-find allows researchers to infer aspects of intentionality and urgency using standard cage conditions and a single sheet of paper. This dramatically increases throughput, preserves the natural environment, and makes behavioral voiding analysis more accessible to a wider range of labs and study designs.

The development of Spoti-find was only possible through the generous collaboration of researchers across institutions who shared pre-existing datasets, allowing us to optimize the tool, test its reproducibility, and compare performance against blinded analyses using Void Whizzard. We view this work as part of a broader effort to support open science. Spoti-find is being released under an open-source license that encourages modification and community-driven improvements, provided that the original work is credited and derivatives are shared non-commercially. As with Void Whizzard, we hope this tool serves as a flexible steppingstone researchers can adapt, refine, and build upon. Given the prevalence of urinary dysfunction across diseases of aging and the nervous system (Hardy and Korstanje 2023), we believe Spoti-find will be broadly useful for identifying and characterizing phenotypes that are often underappreciated in mouse models. Efforts to improve the clinical relevance of behavioral assays -- particularly in areas of urinary function and behavioral control, where cross-species anatomical and neurological differences can obscure interpretation -- are essential for translating preclinical findings into human benefit. By equipping researchers with new ways to analyze, standardize, and quantify VSA data, Spoti-find aims to advance both urologic and broader biomedical research.

These analyses support the utility of Spoti-find’s novel parameters to extract translationally meaningful information from VSA data that has previously been difficult or impossible to quantify. In the clinic, bladder diaries are often used to track symptoms such as frequency, urgency, leakage, and incontinence (Mehta, Geng et al. 2023) -- many of which are behaviorally complex and not captured by void count or volume alone. By incorporating features like circularity, which may reflect whether a voiding event was intentional or passive, and distance from the edge of the cage liner, which may relate to voiding urgency or anxiety-like behavior, Spoti-find allows researchers to approximate aspects of urinary dysfunction that are routinely assessed in human patients. These parameters may be particularly relevant in models of aging or neurodegeneration, where voiding behaviors are altered but not always reflected in total volume voided. As such, Spoti-find enhances the biological resolution and clinical relevance of VSA analysis, enabling richer phenotyping from a simple and scalable assay.

However, this advance comes with clear limitations. Spoti-find is not designed for high-throughput pipelines; its manual interface makes it unsuitable for analyzing thousands of assays efficiently. In such use cases, tools like ML-UrineQuant (Hill, MacIver et al. 2025) may be a better solution. Similarly, while parameters like circularity and edge distance offer novel biological insight, they are inherently more variable and may be affected by user technique, particularly when analyzing small voids with fine structure. The relevance of nanovoids, and in some cases microvoids, remains uncertain, and conclusions drawn from these bins should be interpreted with caution. Nonetheless, the inclusion of these metrics provides a foundation for exploratory phenotyping and future standardization.

The broader utility of Spoti-find will likely become even more apparent as it is applied to additional neurodegenerative disease models, many of which are associated with well-documented urinary phenotypes in humans. Given established bladder dysfunction in models of amyotrophic lateral sclerosis (ALS), Parkinson’s disease, and Alzheimer’s disease, we anticipate that Spoti-find’s novel parameters will provide meaningful insight into voiding behavior across a wide range of neurological conditions. The non-invasive nature of the VSA will permit longitudinal assessments of disease progression overtime without compromising experimental endpoints (i.e., tissue collection, other longitudinal behavior measures) while providing insight into changes along the brain-bladder axis, although physiological assessments (such as pharmacomyography and cystometry) will be required to fully understand the mechanistic changes responsible for altered voiding behavior.

## Methods

### Animals

For the data collected at The Jackson Laboratory, animals were maintained on aspen shavings and given a standard rodent diet (LabDiet 5KOG) and acidified water in a pathogen free room. The room was maintained at 21°C with a 12-hour light/dark cycle (6am to 6pm). All animal experiments were performed in accordance with the National Institutes of Health Guide for the Care and Use of Laboratory Animals (National Research Council) and were approved by The Jackson Laboratory’s Animal Care and Use Committee.

### Void Spot Assay

At The Jackson Laboratory, mice were placed in a cage lined with filter paper (Grade 328 from Alstrom 2381-1417) for 4 hours. After the animal was removed, the paper was dried for a minimum of 24 hours and exposed to UV (details). An image was taken using a Syngene Ingenious bioimaging system (Syngene International).

### Program development

Spotifind was developed in Python. The code and supporting documents, including the installation instructions, user manual, and validation images can be found at https://github.com/TheJacksonLaboratory/Spoti-find. Briefly we use intensity thresholds to identify the paper from background, if the default settings do not work well with the lighting of experimental papers this can be adjusted in the program. Urine spots are identified either using a threshold-based method with the rectangle tool or manually using the polygon tool. Once a spot is identified several measurements are taken. Each spot is counted and classified based on number of pixels compared to a calibration data sheet. Additionally, eccentricity and distance from the center of the spot to nearest edge are calculated.

The data in this paper was analyzed according to the user manual. A combination of rectangle tool and manual polygon spot identification was used. To ensure that users identify spots with similar accuracy and precision, we created a validation data set that is available at https://github.com/TheJacksonLaboratory/Spoti-find and also includes a file with results from three different observers and R code to calculate the interclass correlation coefficient (ICC). Each new user should have over 90% agreement as measured by the ICC before proceeding to experimental papers. We explored the fatigue matrix with 10, 20, 30 and 40 images, Users were given the guidance to score these images taking no breaks until they competed the prescribed number.

### Statistical Analyses

All Statistics were done in R using the open-source packages (DescTools, Lme4 and tidyverse). Anova or linear mixed models were used based on data structure to compare groups with the *aov* function or comparing the fit of linear mixed models using *lme* from the lmer4 package. Inter reader correlation coefficients were obtained using the ICC function.

## Data availability

The datasets used in this study have been deposited at Images.jax.org specific links to each dataset and a brief description of each is below.

**Validation Dataset:** A dataset of 28 Images used to assess inter-reader correlation coefficients and supplied with scores from three people for users to benchmark their spot identification. Images available at: https://images.jax.org/webclient/?show=dataset-5501.

**10-Image Dataset:** A set of ten images from males and females on a C57BL/6J background. Images available at: https://images.jax.org/webclient/?show=dataset-5506.

**20-Image Dataset:** A set of twenty images from males and females on a C57BL/6J background. Images available at: https://images.jax.org/webclient/?show=dataset-5502.

**30-Image Dataset:** A set of thirty images from males and females on a C57BL/6J background. Three readers scored this dataset without breaks. Images available at: https://images.jax.org/webclient/?show=dataset-5503.

**Aging Dataset:** A set of images from 25 young (6-7 weeks) and 25 old (112-113 weeks), male C57BL/6J mice from the Ricke Lab. Images available at: https://images.jax.org/webclient/?show=dataset-5505.

**Disease Dataset:** A set of images from 18 female C57BL/6J mice at 12 weeks old. Control and Cuprizone treatment mice are included papers from the Crocker Lab. Images available at: https://images.jax.org/webclient/?show=dataset-5504

## Code availability

Code for Spoti-find is available on Github (https://github.com/TheJacksonLaboratory/Spoti-find).

## Acknowledgements

We would like to wish Jim Peterson, former computational scientist at The Jackson Laboratory (JAX) a wonderful retirement and thank him for his countless efforts on the initial program development of Spoti-find. We would also like to acknowledge Abigail Brackett, Kevin Koegel, and Samantha Spellacy of JAX for their help with blinded analysis, training protocol development, and inter-user variability assessment. This work was funded through T32AG062409 (CCH) and P30AG038070 (RK).

## Author Contributions

CCH & RK developed the initial idea for this software, including the novel analytical parameters. CCH, SS, and RK wrote the manuscript, with all authors participating in editing and review. SM, NS, and MM developed the necessary coding for the analytical parameters and created the user interface. WR, ZD, and SC provided datasets used in this paper from both published and unpublished sources, including some complementary Void Whizzard datasets. Analyses were primarily conducted by SS. Data interpretation was conducted by CCH, SS, and RK.

## Figure Legends

**Supplemental Table 1.** Inter class correlation to determine agreement between assessments by different observers. Each set was scored by three observers and the inter class correlation (ICC) was calculated. Most values are within the ‘near perfect’ range (>0.81), while ‘micro average volume’, ‘nano average volume’ and ‘micro distance to edge’ are the variables that have correlation coefficients below this range in multiple datasets.

| Phenotype | Validation Set | 10 Images | 20 Images | 30 Images |
| --- | --- | --- | --- | --- |
| Paper Area | 1.000 | 0.983 | 0.983 | 1.000 |
| Primary Count | 0.938 | 0.948 | 0.972 | 0.930 |
| Micro Count | 0.918 | 0.878 | 0.949 | 0.856 |
| Nano Count | 0.968 | 0.893 | 0.951 | 0.926 |
| Total Count | 0.942 | 0.939 | 0.959 | 0.888 |
| Primary Volume | 0.972 | 0.877 | 0.861 | 0.981 |
| Micro Volume | 0.876 | 0.889 | 0.886 | 0.695 |
| Nano Volume | 0.969 | 0.909 | 0.972 | 0.960 |
| Total Volume | 0.972 | 0.875 | 0.862 | 0.981 |
| Primary Average Volume | 0.918 | 0.810 | **0.569** | 0.925 |
| Micro Average Volume | **0.639** | 0.918 | **0.319** | 0.940 |
| Nano Average Volume | **0.436** | **0.720** | **0.736** | **0.755** |
| Total Average Volume | 0.946 | 0.877 | 0.929 | 0.944 |
| Primary Distance to Edge | 0.983 | 0.974 | 0.984 | 0.979 |
| Micro Distance to Edge | **0.781** | **0.467** | **0.788** | 0.964 |
| Nano Distance to Edge | 0.917 | 0.900 | 0.928 | 0.810 |
| Total Distance to Edge | 0.908 | 0.967 | 0.937 | 0.896 |

